# SARS-CoV-2 variants with mutations at the S1/S2 cleavage site are generated *in vitro* during propagation in TMPRSS2-deficient cells

**DOI:** 10.1101/2020.08.28.271163

**Authors:** Michihito Sasaki, Kentaro Uemura, Akihiko Sato, Shinsuke Toba, Takao Sanaki, Katsumi Maenaka, William W. Hall, Yasuko Orba, Hirofumi Sawa

## Abstract

The spike (S) protein of Severe Acute Respiratory Syndrome-Coronavirus-2 (SARS-CoV-2) binds to a host cell receptor which facilitates viral entry. A polybasic motif detected at the cleavage site of the S protein has been shown to broaden the cell tropism and transmissibility of the virus. Here we examine the properties of SARS-CoV-2 variants with mutations at the S protein cleavage site that undergo inefficient proteolytic cleavage. Virus variants with S gene mutations generated smaller plaques and exhibited a more limited range of cell tropism compared to the wild-type strain. These alterations were shown to result from their inability to utilize the entry pathway involving direct fusion mediated by the host type II transmembrane serine protease, TMPRSS2. Notably, viruses with S gene mutations emerged rapidly and became the dominant SARS-CoV-2 variants in TMPRSS2-deficient cells including Vero cells. Our study demonstrated that the S protein polybasic cleavage motif is a critical factor underlying SARS-CoV-2 entry and cell tropism. As such, researchers should be alert to the possibility of *de novo* S gene mutations emerging in tissue-culture propagated virus strains.

## Introduction

The World Health Organization has declared disease (COVID-19) due to infection with Severe Acute Respiratory Syndrome-Coronavirus-2 (SARS-CoV-2) as pandemic. As of August 6, 2020, more than 18 million confirmed cases and 720,000 fatalities have been reported worldwide [1]. The SARS-CoV-2 virion includes four structural elements identified as the spike (S), envelope (E), membrane (M), and nucleocapsid (N) proteins [2, 3]. The S protein forms a homotrimer on the virion surface and triggers viral entry into target cells via interactions between its receptor binding domain and the specific host receptor, angiotensin-converting enzyme 2 (ACE2) [4, 5]. Two pathways have been proposed for virion entry into cells; these include direct fusion at the plasma membrane mediated by the host type II transmembrane serine protease, TMPRSS2, and endocytic entry that relies on the actions of the lysosomal protease, cathepsin [6]. SARS-CoV-2 utilizes both entry pathways for infection of cells that express TMPRSS2, while TMPRSS2-deficient cells permit viral entry exclusively *via* the cathepsin-dependent endosome pathway [6].

The S protein of SARS-CoV-2 includes a discriminative polybasic cleavage motif (RRAR) at the S1/S2 cleavage site which is not present in the S proteins of related coronaviruses, including human SARS-CoV (Figs. 1a and 1b) [7, 8]. This polybasic motif has been shown to facilitate the cleavage of nascent S protein into S1 and S2 subunits by the host furin protease; cleavage is critical for formation of cell syncytia and for efficient entry of vesicular stomatitis virus (VSV) pseudotyped with SARS-CoV-2 S protein into host target cells [7]. Consequently, the polybasic cleavage motif of the SARS-CoV-2 S protein has emerged as a feature of significant interest and importance.

**Fig. 1.**
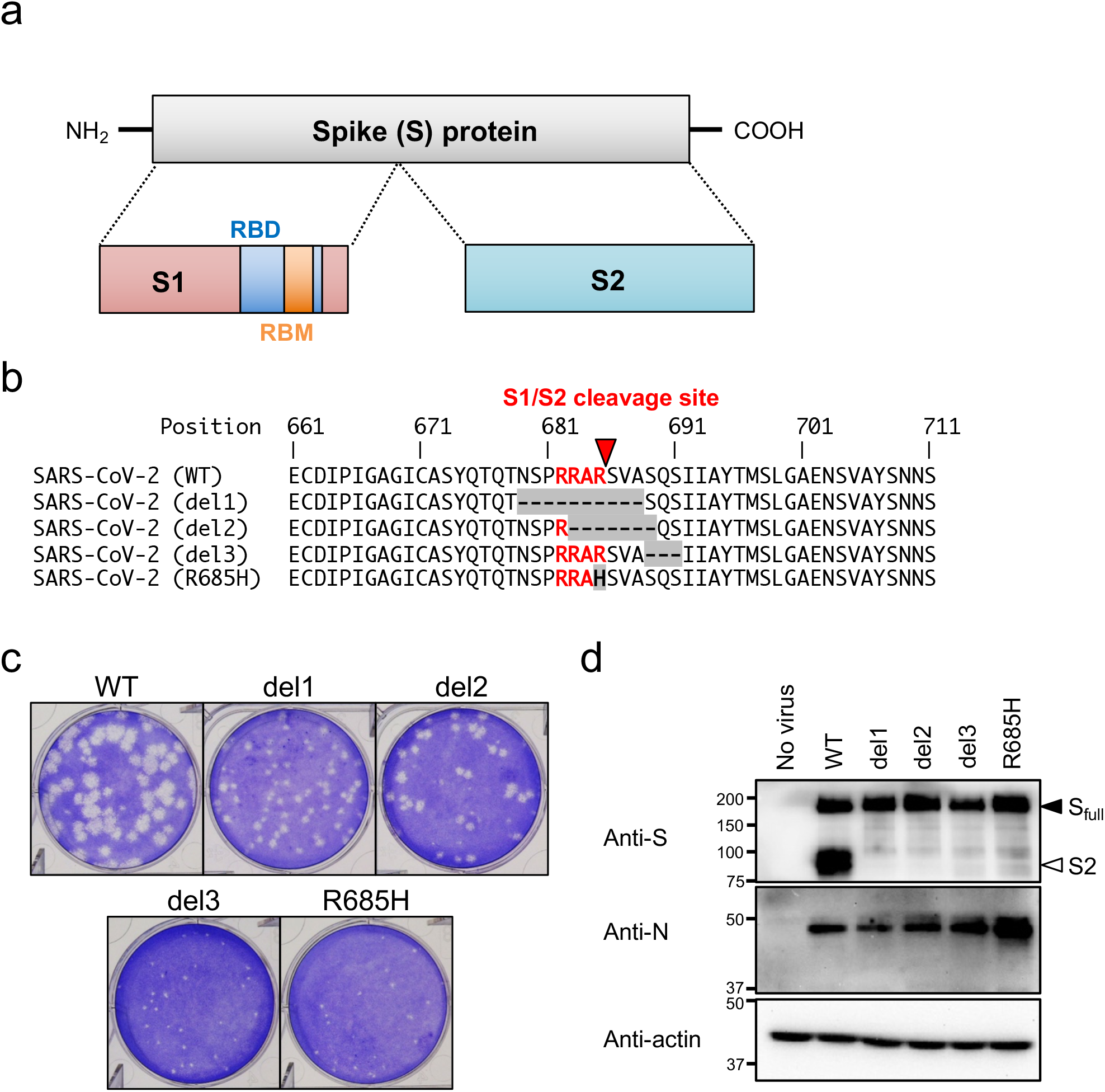
Isolation of SARS-CoV-2 S gene mutants. (a) Schematic representation of SARS-CoV-2 S protein. Full-length S protein is cleaved into S1 and S2 proteins at the S1/S2 cleavage site. Functional domains (RBD, receptor binding domain; RBM, receptor binding motif) are highlighted. (b) Multiple amino acid sequence alignments focused on the S1/S2 cleavage site of wild type (WT) and isolated mutant viruses (del1, del2, del3 and R685H). Amino acid substitutions and deletions are shown as gray boxes, and the polybasic cleavage motif (RARR) at the S1/S2 cleavage site is highlighted in red. A red arrowhead indicates S1/S2 cleavage site. (c) Plaque formation for SARS-CoV-2 WT and isolated mutants grown in Vero-TMPRSS2 cells on 6 well plates. (d) Detection of virus S and N proteins in Vero-TMPRSS2 cells infected with WT or isolated mutants. The full-length S and cleaved S2 proteins are indicated by closed and open arrowheads, respectively.

In this study, we have examined the properties of SARS-CoV-2 variants which have developed mutations at the S protein cleavage site during routine passage in cell culture. The variants with S gene mutations were unable to utilize the virus entry pathway mediated by TMPRSS2 and exhibited a more limited range of cell tropism compared to the wild-type strain. Notably, SARS-CoV-2 variants with S gene mutations emerged rapidly and became the dominant in the virus population after two passages in Vero cells, which are common cell targets used for the study of SARS-CoV-2.

## Results

### Isolation of SARS-CoV-2 S gene mutants

Vero cells express ACE2 under homeostatic conditions and are susceptible to infection with SARS-CoV-2 [9, 10] and this cell line is used widely for isolation and propagation of SARS-CoV-2. Currently, Vero cells engineered to maintain stable expression of TMPRSS2 (Vero-TMPRSS2) are highly susceptible to SARS-CoV-2 infection [11]. After several passages, we detected variant viruses with mutations at the S1/S2 cleavage site in the S gene of SARS-CoV-2 (S gene mutants) when virions were propagated in Vero cells. In contrast, these were not detected in viruses maintained in Vero-TMPRSS2. In order to characterize the properties of the S gene mutant viruses, we isolated four variant clones from progeny virus pools by limiting dilution. Nucleotide sequence analysis revealed that three of the isolated virus clones (del1, del2, and del3) had in-frame deletions; the fourth variant (R685H) had a single nucleotide substitution at the S1/S2 cleavage site (Fig. S1). Multiple alignment analysis revealed that del1 and del2 had deletions of 10 and 7 amino acid residues, respectively, and these included the polybasic cleavage motif (RRAR) at the S protein S1/S2 cleavage site. In contrast, del3 had a deletion of 3 amino acids at a point immediately downstream of RRAR motif (Fig. 1b). In R685H, the RRAR motif was substituted with RRAH (Fig. 1b).

At 3 days post infection, viruses with S gene mutations generated smaller plaques on Vero-TMPRSS2 cells compared with the parent (WT) strain (Fig. 1c). Earlier reports revealed that the introduction of mutations at the polybasic cleavage motif prevented effective cleavage of the SARS-CoV-2 S protein [4, 7]. As all mutants isolated in our study included deletions or substitutions within or near the polybasic cleavage motif, we evaluated the extent of cleavage of the nascent S protein by immunoblotting with an anti-S monoclonal antibody which detects both full-length S protein and its S2 cleavage product. Vero-TMPRSS2 cells were susceptible to the infection with WT and S gene mutant viruses and infection was associated with comparable levels of nascent viral N and full-length S proteins (Fig. 1d). However, S2 protein was detected only in cells infected with WT virus. Notably, while the polybasic RRAR motif of del3 remained intact, S protein cleavage in this mutant variant was impaired in a manner similar to that observed among the other S gene mutants. Taken together, these results suggested that amino acid deletion or substitution in or near the SARS-CoV-2 S1/S2 cleavage site prevented S protein cleavage and resulted in decreased plaque sizes.

### Cell tropism of S gene mutants

We then investigated the tropism and growth of S gene mutant viruses in different cell lines. Immunofluorescence analysis revealed that Vero and Vero-TMPRSS2 cells were both highly susceptible to infection with WT and all S gene mutant viruses (Fig. 2a). However, formation of characteristic cell syncytia was observed only in Vero-TMPRSS2 cells infected with WT virus (white arrowheads in Fig. 2a). The human lung epithelial Calu-3 and colon epithelial Caco-2 cell lines are known to be susceptible to SARS-CoV-2 infection [10]. However, compared to WT virus, few to no Calu-3 or Caco-2 cells were susceptible to infection with any of the S gene mutant viruses (Fig. 2a). Likewise, although the virus titers of S gene mutants generated in infected Vero and Vero-TMPRSS2 cells were similar to those of the WT virus, titers of S gene mutants were significantly lower those of the WT virus in Calu-3 and Caco-2 cells, and these results correlate with the findings of immunofluorescence analysis (Fig. 2b). Notably, the virus titers from Calu-3 cells inoculated with del1 and del2 were under the limits of detection at all time points. These results indicate that both Calu-3 and Caco-2 cells were less susceptible to infection with S gene mutants compared to WT virus. This was intriguing, given that both Calu-3 and Caco-2 cells express endogenous ACE2 [9, 12] and TMPRSS2 [13, 14].

**Fig. 2.**
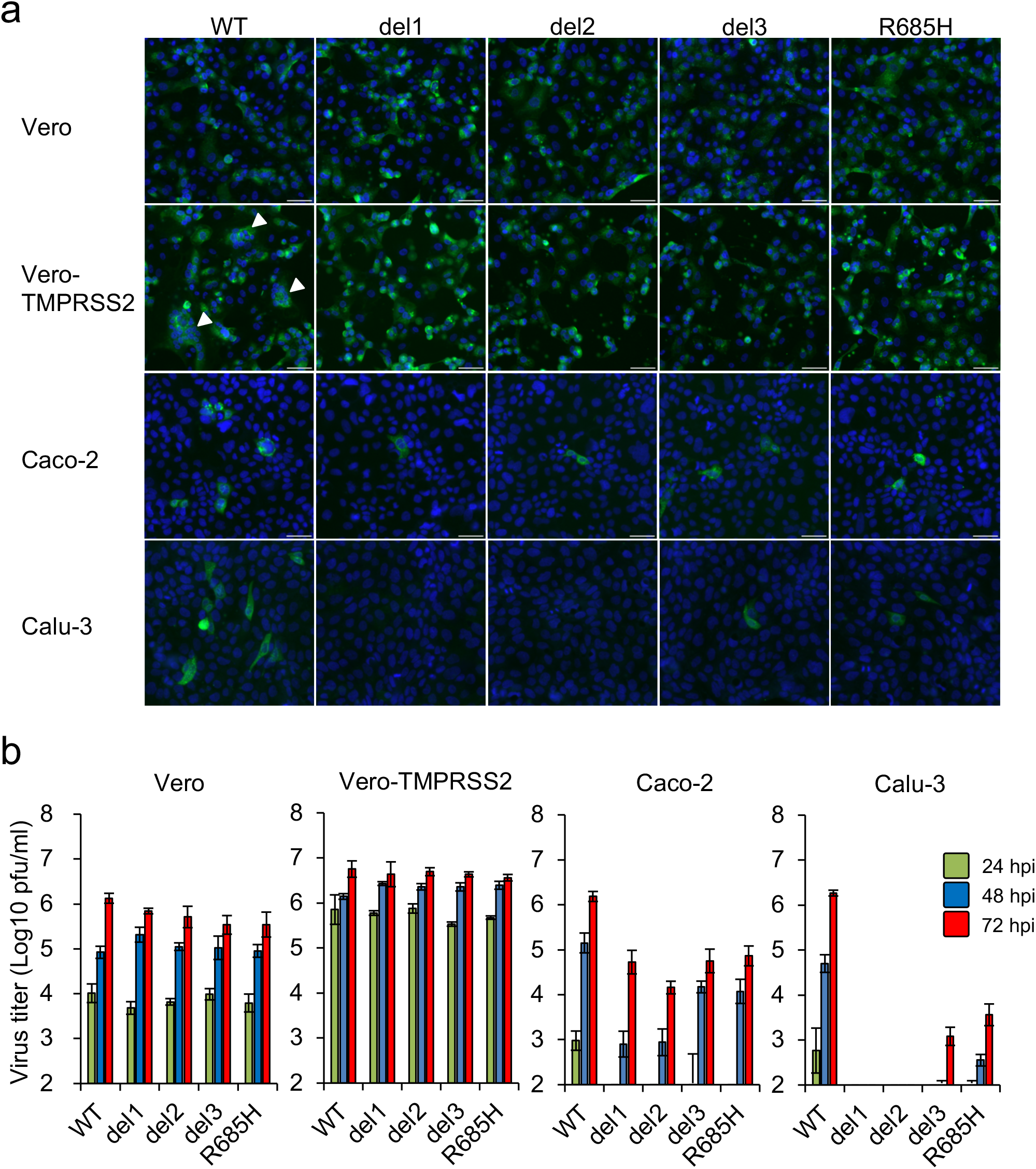
Infection and growth of SARS-CoV-2 S gene mutants in different cell lines. (a) Vero, Vero-TMPRSS2, Caco-2 and Calu-3 cells were inoculated with SARS-CoV-2 WT or S gene mutants at a multiplicity of infection (MOI) of 1. At 24 h post infection, cells were stained with anti-SARS-CoV-2 S antibody (green) and Hoechst 33342 nuclear dye (blue); scale bars, 50 μm. White arrowheads indicate cell syncytia. (b) Cells were inoculated with SARS-CoV-2 WT or S gene mutants at an MOI of 0.1. Culture supernatants were harvested at 24, 48 and 72 h after inoculation. Virus titration was performed by plaque assay. The values shown are mean ± standard deviation (SD) of triplicate samples.

Human kidney 293T cells lack ACE2 expression and its exogenous introduction confers susceptibility to SARS-CoV-2 infection [6, 15]. To examine the receptor usage by the S gene mutants, 293T cells at baseline and 293T cells stably expressing human ACE2 (293T-ACE2) cells were inoculated with WT virus or S gene mutants. Heterologous expression of ACE2 resulted in a marked increase susceptibility to infection with the WT and the S gene mutant viruses (Figs. 3a and 3b) and titers of S gene mutants were equivalent or higher than that those of the WT virus infection of 293T-ACE2 cells (Fig. 3b). These results suggest that ACE2 facilitates cell entry and infection of both S gene mutants and the WT virus.

**Fig. 3.**
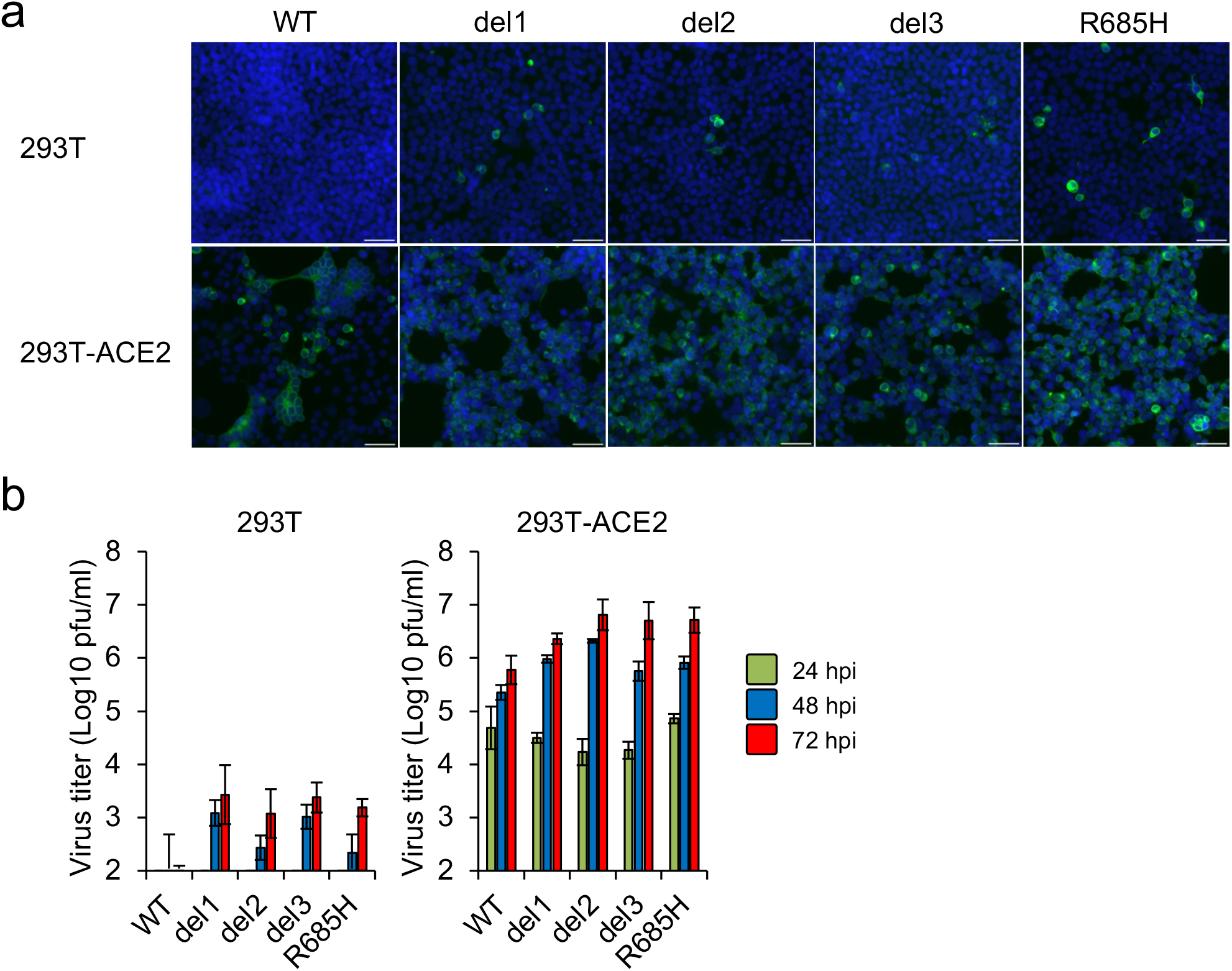
Infection and growth of SARS-CoV-2 S gene mutants in 293T and 293T-ACE2 cells. (a) 293T and 293T-ACE2 cells were inoculated with SARS-CoV-2 WT or S gene mutants at an MOI of 1. At 24 h after inoculation, cells were stained with anti-SARS-CoV-2 S antibody (green) and Hoechst 33342 nuclear dye (blue); scale bars, 50 μm. (b) Cells were infected with SARS-CoV-2 WT or S gene mutants at an MOI of 0.1. Culture supernatants were harvested at 24, 48 and 72 h after inoculation and titration of infectious virus was determined by plaque assay. The values shown are means ± SD of triplicate samples.

### Effect of chemical inhibitors on entry of S gene mutants

Given that there were no differences with respect to ACE2 usage, we next examined whether S gene mutants utilize one or both cellular entry pathways. These experiments were performed with camostat mesylate, an inhibitor of TMPRSS2, and E-64d, an inhibitor of cathepsin B/L. These agents are known to inhibit cellular entry of coronaviruses, including SARS-CoV-2 [6, 16]. Camostat inhibited WT SARS-CoV-2 entry into Vero-TMPRSS2 cells; however E-64d had no impact on this process (Fig. 4a). These findings were consistent with a previous study that reported that TMPRSS2-mediated entry was the dominant pathway employed by SARS-CoV-2 in TMPRSS2-expressing cells [6]. In contrast, camostat had no impact on entry of the S gene mutant, del2, into Vero-TMPRSS2 cells but E-64d treatment resulted in a dose-dependent decrease in del2 entry (Fig. 4a). These results suggested that S gene mutant, del2, can enter Vero-TMPRSS2 cells via cathepsin-dependent endocytosis but not the TMPRSS2-mediated fusion pathway. Parental Vero cells that do not express TMPRSS2 were inoculated with S gene mutant viruses in the presence of camostat and/or E-64d. Addition of E-64d inhibited the entry of all S gene mutants into both Vero-TMPRSS2 and parent Vero cells; by contrast, camostat had no impact on S gene mutant entry into these target cells (Fig. 4b). These results suggested that, in contrast to WT virus, S gene mutants enter into cells via cathepsin-dependent endocytosis only, regardless of the presence or absence of TMPRSS2.

**Fig. 4.**
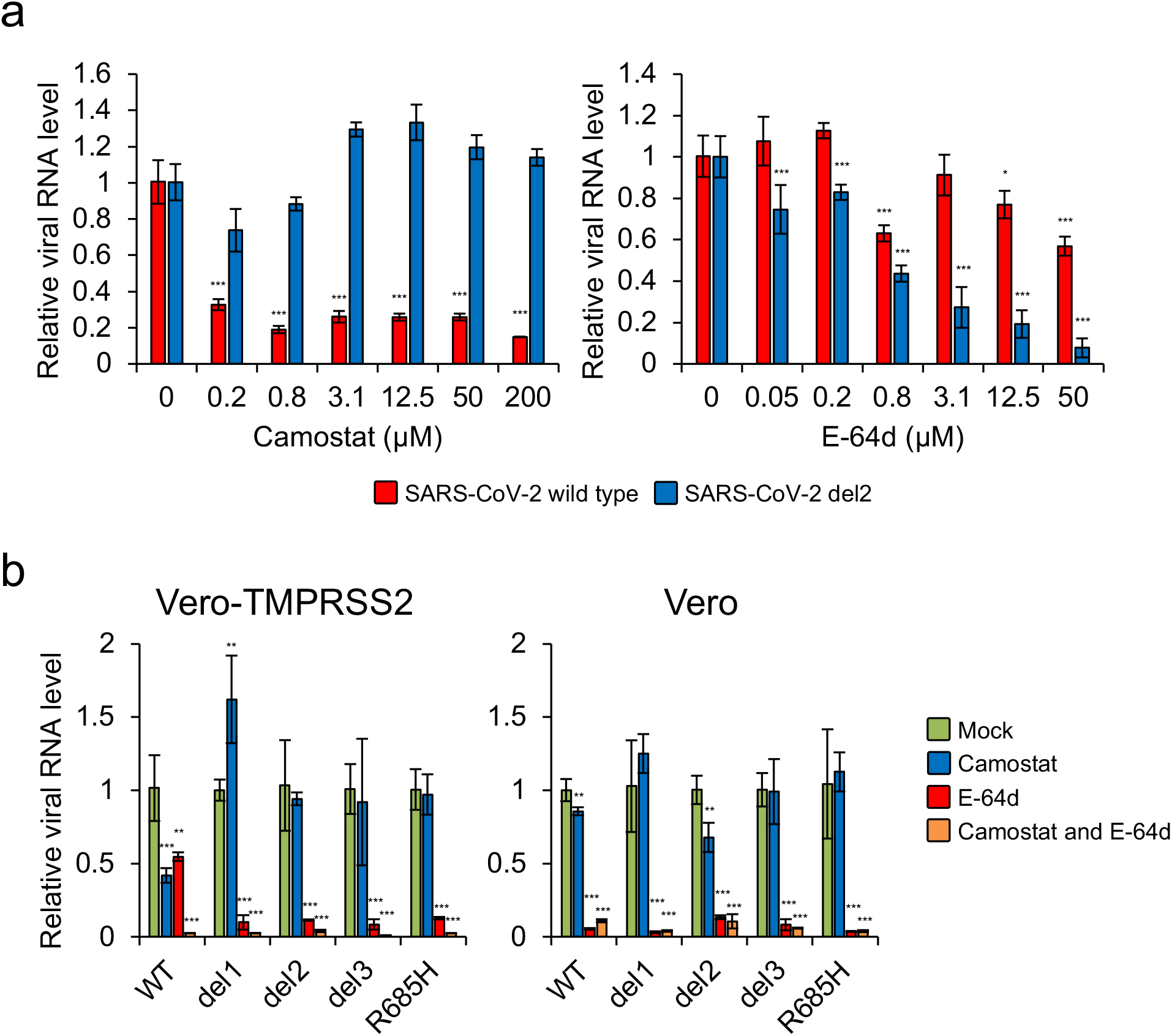
Impact of biochemical inhibitors on cellular entry of SARS-CoV-2 S gene mutants. (a) Vero-TMPRSS2 cells were infected with SARS-CoV-2 WT or del2 mutant in the presence of varying concentrations of the TMPRSS2 inhibitor, camostat, or the cathepsin B/L inhibitor, E-64d, for 1h. At 6 h post-inoculation, the relative levels of viral N protein RNA were evaluated quantitatively by qRT-PCR. (b) Vero-TMPRSS2 and Vero cells were infected with SARS-CoV-2 WT or S gene mutants in the presence of 50 μM camostat and/or 25 μM E-64d for 1 h. At 6 h post-inoculation, the relative levels of viral N protein RNA were quantified by qRT-PCR. Cellular β-actin mRNA levels were used as reference controls. The values shown are mean ± SD of triplicate samples. One-way analysis of variance with Dunnett’s test was used to determine the statistical significance between the responses to treatment with inhibitors and the no-treatment controls; \**p* < 0.05, \*\**p* < 0.01, \*\*\**p* < 0.001.

Because WT virus and S gene mutants showed different sensitivities to the treatment with camostat, an agent currently under exploration as a candidate antiviral for clinical use [17], we also examined the impact of other antiviral agents including nafamostat (a TMPRSS2 inhibitor) [18, 19] and remdesivir (a nucleotide analog) [20, 21]. Antiviral effects in Vero-TMPRSS2 cells were estimated by a cell viability assay based on the generation of cytopathic effects [22]. Consistent with previous studies [18, 19], nafamostat showed higher antiviral efficacy against WT virus than was observed in response to camostat; however, nafamostat had no antiviral activity against the S gene mutants (Table 1). In contrast, remdesivir inhibited infection of both WT and S gene mutants with similar EC_50_ values (Table 1). These results indicated that S gene mutants are resistant to the treatment with TMPRRSS2 inhibitors, but are sensitive to antivirals that target post entry processes.

**Table 1.**
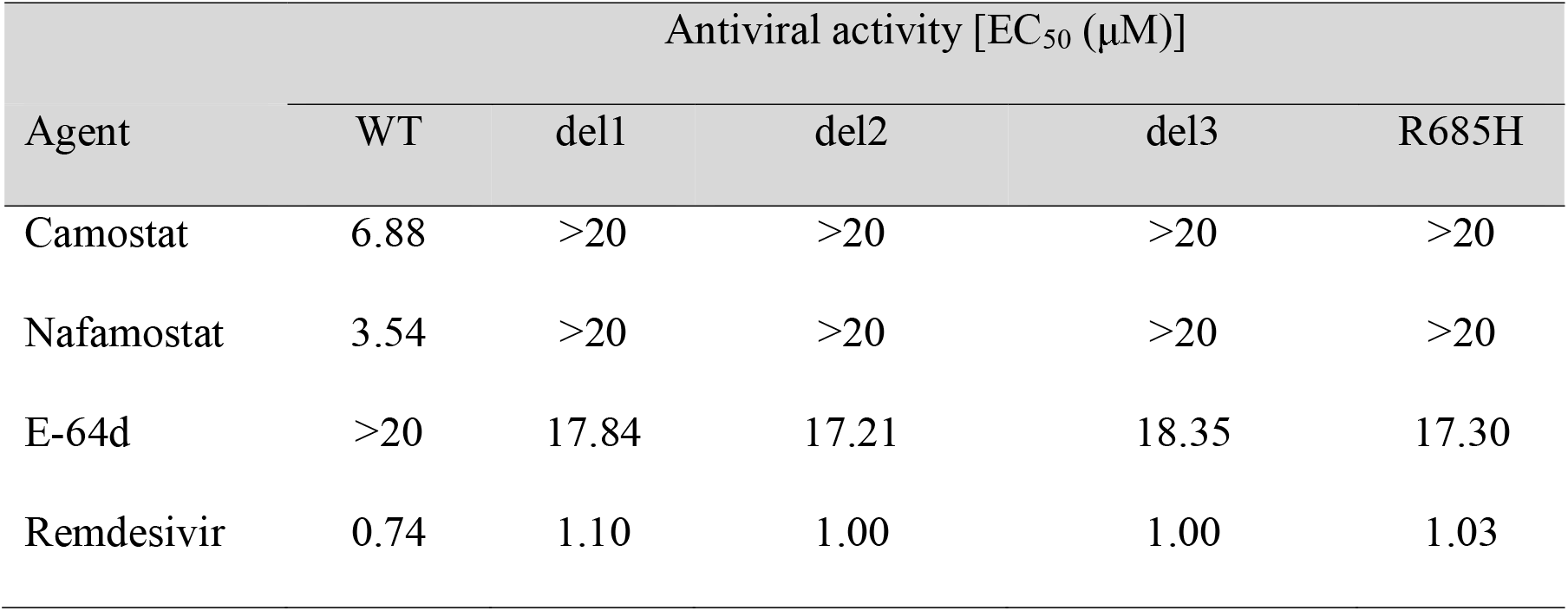
Impact of antivirals against WT and S gene mutants of SARS-CoV-2 in Vero-TMPRSS2 cells.

| Agent | Antiviral activity [EC<sub>50</sub> (μM)] |  |  |  |  |
| --- | --- | --- | --- | --- | --- |
|  | WT | del1 | del2 | del3 | R685H |
| Camostat | 6.88 | >20 | >20 | >20 | >20 |
| Nafamostat | 3.54 | >20 | >20 | >20 | >20 |
| E-64d | >20 | 17.84 | 17.21 | 18.35 | 17.30 |
| Remdesivir | 0.74 | 1.10 | 1.00 | 1.00 | 1.03 |

### Frequencies of S gene mutants in SARS-CoV-2 propagation

In an effort to understand the selection mechanisms underlying the generation of these mutant variants, we estimated the frequency of S gene mutants in virus population of SARS-CoV-2 that had undergone serial passage in cultured cells. SARS-CoV-2 from an original virus stock was underwent passage once (P1) to four times (P4) in Vero or up to eight times (P8) in Vero-TMPRSS2. Nucleotide sequence heterogeneity at the S1/S2 cleavage site was determined by deep sequencing and variant call analysis. More than one million sequence reads from each passaged sample were mapped onto the S1/S2 cleavage site and analyzed for sequence variation. No sequence variants were observed in virus populations until P8 in Vero-TMPRSS2 (Fig. 5a). In contrast, nucleotide sequence deletions around the S1/S2 cleavage site corresponding to del1 and del2 mutants were observed in all three biological replicates of SARS-CoV-2 populations passaged in Vero cells (Fig. 5a). Notably, WT nucleotide sequences were detected in fewer than 20% of the isolates evaluated at P2 and the WT was completely replaced with S gene mutants at P4 in Vero cells. These results indicated that SARS-CoV-2 propagation in Vero cells results in a profound selection favoring the S gene mutants. S gene mutants del3 and R685H were not identified in the virus populations from P1 to P4. An additional variant del4 with a deletion of 5 amino acids at a point immediately upstream of the RRAR motif (Figs. S2a and S2b), was detected as a minor variant in sample #1 at P2. These results suggest that these specific mutations occur only at low frequency.

**Fig. 5.**
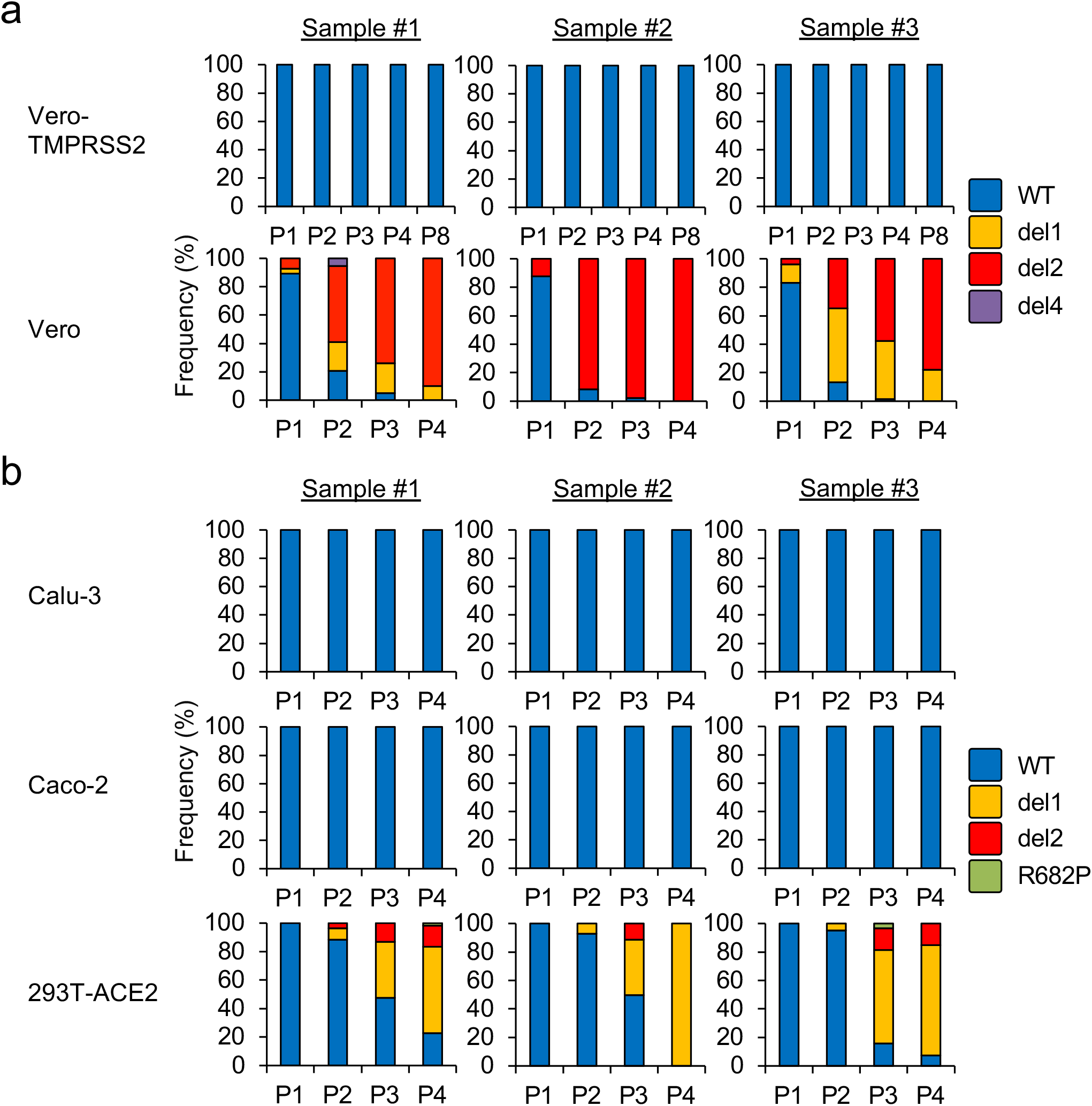
Frequencies of S gene mutants detected during SARS-CoV-2 propagation. SARS-CoV-2 was serially passaged in (a) Vero-TMPRSS2, Vero cells or (b) Calu-3, Caco-2 and 293T-ACE2 cells, each with three biological replicates. Nucleotide sequence diversity at viral S1/S2 cleavage site was determined by deep-sequencing.

We then determined the frequency of S gene mutants in virus populations passaged in Calu-3 and Caco-2, which are cells that endogenously express TMPRSS2 [13, 14], and also in 293T-ACE2 that do not express TMPRSS2 [6]. No S gene mutants were identified in SARS-CoV-2 passaged in Calu-3 and Caco-2 until P4; by contrast, S gene mutants emerged at P2 in 293T-ACE2 cells (Fig. 5b). We also identified an additional variant R682P carrying a single amino acid substitution at the RRAR motif (Figs. S2a and S2b) at P3 and P4 in 293T-ACE2 cells (Fig. 5b). Taken together, these results suggest a strong association between TMPRSS2 deficiency and the emergence of S gene mutants.

Trypsin is a serine protease that is typically added to culture medium to induce cleavage and activation of viral proteins, including the hemagglutinin (HA) protein of influenza virus and the fusion (F) protein of paramyxovirus to promote growth in TMPRSS2-deficient cells [23]. Recent studies report that trypsin treatment activates SARS-CoV-2 S protein and induces syncytia formation in cells that transiently express the virus S protein [7, 24]. As such, we examined whether exogenously added trypsin could compensate for TMPRSS2 deficiency and thus inhibit the emergence of S gene mutants during SARS-CoV-2 propagation in Vero cells. Deep sequencing analysis revealed that S gene mutants did emerge and accounted for the majority of the virus population after P2 in Vero cells cultured in serum-free medium with added trypsin (Fig. S3); these results indicate that exogenous protease activity cannot replace TMPRSS2 in its role in promoting SARS-CoV-2 replication.

## Discussion

In this study, we isolated S gene mutants from SARS-CoV-2 WK-521, a strain isolated from a clinical case in Japan [11], via serial passage in Vero cells. Other studies have reported viruses with S gene mutations, including amino acid deletions and substitution at the S1/S2 cleavage site from clinical isolates in Australia [20], China [25], and England [26] and these emerged during cultivation in Vero cells or in its derivative, Vero/hSLAM, which are cells that do not express TMPRSS2. However, the virological properties of the S gene mutants remains to be poorly investigated. Our deep sequencing analysis revealed that S gene mutants emerged at P1 and rapidly became the dominant variant within the virus populations that emerged from Vero cell passage. Taken together, our findings indicate that replication of SARS-CoV-2 in TMPRSS2-deficient Vero cells results in the selection of S gene mutants; as such, passage in this cell line is technically inappropriate, as it becomes difficult to impossible to maintain SARS-CoV-2 with the S1/S2 cleavage site in its intact form.

The present study characterized S gene mutants as SARS-CoV-2 variants that generate small plaques and that have a narrow range of cell tropism. The phenotypic alterations of S gene mutants might be explained by noting that the S gene mutants were unable enter target cells via direct fusion mediated by TMPRSS2. Indeed, a previous study demonstrated that the polybasic cleavage motif at the S1/S2 cleavage site was indispensable for the entry of VSV-pseudotyped viruses into Calu-3 cells that expressed TMPRSS2 [7]. Further studies using infectious S gene mutants will provide new insights into the role of the polybasic amino acid motif at the S1/S2 cleavage site with respect to both SARS-CoV-2 infection and its pathogenicity.

At this time, many studies are conducted using SARS-CoV-2 propagated in Vero cells. Considering the very real possibility that these virus stocks will accumulate S gene mutations, researchers must pay careful attention to the passage history of any working stocks of SARS-CoV-2. Moreover, we must be very objective when interpreting the results from studies using Vero-passaged virus, especially those focused on S protein cleavage, virus entry and on cell tropism of SARS-CoV-2.

## Methods

### Cells

Calu-3 (ATCC) were maintained in Eagle’s Minimum Essential Medium (MEM) supplemented with 10% fetal bovine serum (FBS). Caco-2 (RIKEN BRC) cells were maintained in Eagle’s MEM supplemented with 10% FBS and non-essential amino acids. Vero E6 (ATCC) and 293T (JCRB cell bank) cells were maintained in Dulbecco’s Modified Eagle’s Medium (DMEM) supplemented with 10% FBS. All cells were incubated at 37°C with 5% CO_2_.

### Generation of TMPRSS2- and ACE2-expressing cells

Human TMPRSS2 and ACE2 genes were cloned into the self-inactivating lentiviral vector plasmids, CSII-CMV-MCS-IRES2-Bsd (RIKEN BRC) and pLVSIN-CMV Pur (Takara Bio), respectively. The resulting constructs were named CSII-CMV-TMPRSS2-IRES2-Bsd and pLVSIN-CMV-ACE2-Pur. For lentiviral vector preparation, 293T cells were co-transfected with the lentiviral vector plasmid and Lentiviral High Titer Packaging Mix (Takara Bio). The culture supernatants containing lentiviral vectors were used to inoculate target cells. Vero cells stably expressing TMPRSS2 (Vero-TMPRSS2) and 293T stably expressing ACE2 (293T-ACE2) were selected in the presence of blasticidin S or puromycin.

### Viruses

SARS-CoV-2 WK-521 strain was provided by Dr. Shimojima (National Institute of Infectious Diseases, Japan); the original stock of this virus (wild type, WT) was prepared by inoculation of Vero-TMPRSS2 cells with Mynox mycoplasma elimination reagent (Minerva Biolabs) [11]. After several passages of SARS-CoV-2 in Vero cells, the S mutant viruses in the culture supernatant were cloned by limiting dilution and propagated in Vero cells. The nucleotide sequences of S genes of all working stocks were confirmed by RT-PCR and direct sequencing methods.

### Plaque assay

Monolayers of Vero-TMPRSS2 were inoculated with serial dilutions of either WT or S mutants of SARS-CoV-2 for 1h at 37°C. The cells were then overlaid with DMEM containing 0.5% Bacto Agar (Becton Dickinson). At 3 days post-inoculation, cells were fixed with 3.7% buffered formaldehyde and stained with 1% crystal violet.

### Immunoblotting

Vero-TMPRSS2 cells were infected with either WT or S mutants of SARS-CoV-2 at a multiplicity of infection (MOI) of 0.1. After 24 h, infected cells were lysed in lysis buffer (1% NP-40, 20 mM Tris-HCl [pH 7.5], 150 mM NaCl, 5 mM EDTA) supplemented with cOmplete ULTRA protease inhibitor cocktail (Roche Diagnostics). Proteins in each lysate were resolved by SDS-PAGE and transferred onto Immobilon-P PVDF membranes (Merck). The blots were incubated with the following primary antibodies: anti-SARS-CoV-2 N or anti-SARS-CoV-2 S (GTX632269, GTX632604, GeneTex). HRP-conjugated anti-β-actin antibody (PM053-7, MBL) was used to detect the loading control. Immune complexes were detected using HRP-conjugated secondary antibodies and the Immobilon Western Chemiluminescent HRP Substrate (Merck).

### Indirect immunofluorescence assay

Cells were infected with either WT or S mutants of SARS-CoV-2 at an MOI of 1. After 24 h, cells were fixed with 3.7% buffered formaldehyde, permeabilized with ice-cold methanol, and incubated with anti-SARS-CoV-2 S antibody (GTX632604, GeneTex). Alexa Fluor Plus 488-conjugated anti-mouse IgG antibody (Invitrogen; Thermo Fisher Scientific) was used as the secondary antibody. Nuclei were stained with Hoechst 33342 (Invitrogen). Fluorescent images were captured using a fluorescence microscope (IX73, Olympus).

### Multi-cycle growth of SARS-CoV-2

Cells were inoculated with either WT or S mutants of SARS-CoV-2 at an MOI of 0.1. After 1 h of incubation, cells were washed twice with phosphate-buffered saline (PBS) and cultured in fresh medium with 2% FBS. The culture supernatants were harvested at 24, 48, and 72 h after inoculation. Virus titers were evaluated by plaque assay.

### Virus infection assay with biochemical inhibitors

Cells were infected with either WT or S mutants of SARS-CoV-2 at an MOI of 0.1 in the presence of 50 μM camostat mesylate (FUJIFILM Wako Pure chemical) and/or 25 μM E-64d (Abcam) for 1 h. Cells were then washed twice with PBS and cultured in fresh culture medium with 2% FBS. At 6 h after inoculation, total RNA was extracted from cells with Direct-zol-96 RNA Kit or Direct-zol RNA MiniPrep Kit (Zymo Research). RNA samples were subjected to qRT-PCR analysis using AgPath-ID One-Step RT-PCR Kit (Applied Biosystems; Thermo Fisher Scientific). The primer and probe sequences targeting SARS-CoV-2 N gene included: 5′–CACATTGGCACCCGCAATC–3′, 5′–GAGGAACGAGAAGAGGCTTG–3′, and 5′-FAM–ACTTCCTCA/ZEN/AGGAACAACATTGCCA/–IBFQ–3′ [27]. The primer and probe sequences for nonhuman primate *β-actin* were as described previously [28].

### Cytopathic effect-based cell viability assays

The MTT (3-[4,5-dimethyl-2-thiazolyl]-2,5-diphenyl-2H-tetrazolium bromide) (Nacalai Tesque) assay was performed to evaluate cell viability following viral infection according to methods previously described [29]. Camostat, E-64d, nafamostat (FUJIFILM Wako Pure chemical) and remdesivir (MedChemExpress) were serially diluted 2-fold increments in duplicates and plated on 96-well microplates in MEM containing 2% FBS. Vero-TMPRSS2 were infected with either WT or S mutants of SARS-CoV-2 at 4-10 TCID_50_ and added to the plates. Plates were incubated at for 3 days, and CPE was determined for calculation of 50% endpoints using MTT assay. The concentration achieving 50% inhibition of cell viability (effective concentration; EC_50_) was calculated.

### Deep sequencing of the S gene of passaged SARS-CoV-2 virions

The original stock of SARS-CoV-2 strain WK-521 was serially passaged in Vero, Vero-TMPRSS2, Calu-3, Caco-2, and 293T-ACE2 cells in complete culture medium or (for Vero) in serum free DMEM supplemented with 0.5 μg/ml trypsin (Gibco); three biological replicates were included for each of the cell lines. Virus propagation was performed in 12-well plates; 20 μl (2%) of culture supernatant collected at day 3 post infection was used to inoculate naïve cells for virus passage. RNA was extracted from the culture supernatant after each passage using a High Pure Viral RNA kit (Roche Diagnostics). For deep sequencing of the S1/S2 cleavage site of the viral S gene, amplicon sequence libraries were fused with Ion A and the Ion Express barcode sequence at the 5′-region; truncated P1 adapters at 3′-region were generated by nested RT-PCR with a fusion method according to the manufacturer’s instructions (Fusion methods from Ion Torrent; Thermo Fisher Scientific). Information on the fusion PCR primers is available upon request. For deep sequencing, emulsion PCR was performed with the Ion PI Hi-Q OT2 200 kit (Ion Torrent). Sequencing was performed using the Ion PI Hi-Q Sequencing 200 kit, the Ion PI Chip Kit v3 and the Ion Proton sequencer (Ion Torrent). After mapping the reads to the reference sequence (GenBank accession no. LC522975), sequence variants including deletions, insertions, and substitutions were identified using the Torrent Variant Caller plugin with indel minimum allele frequency cutoff of 0.02 (Ion Torrent).

### Statistical analysis

One-way analysis of variance with Dunnett’s test was employed to determine statistical significance.

## Acknowledgments

We thank Dr. Shimojima at National Institute of Infectious Diseases, Japan for providing SARS-CoV-2 WK-521 strain and Dr. Miyoshi at RIKEN BRC, Japan for providing lentiviral vector plasmid CSII-CMV-MCS-IRES2-Bsd.

---

**Fig. S1.**
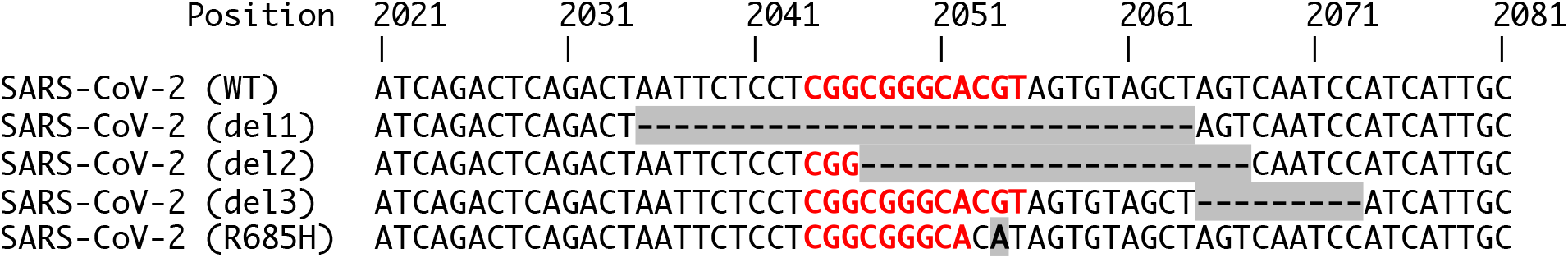
Multiple nucleotide sequence alignment of S1/S2 cleavage site of wild type and isolated SARS-CoV-2 mutants. Nucleotide substitutions and deletions are shown as gray boxes. Sequence encoding the polybasic cleavage motif (RARR) at the S1/S2 cleavage site is highlighted in red.

**Fig. S2.**
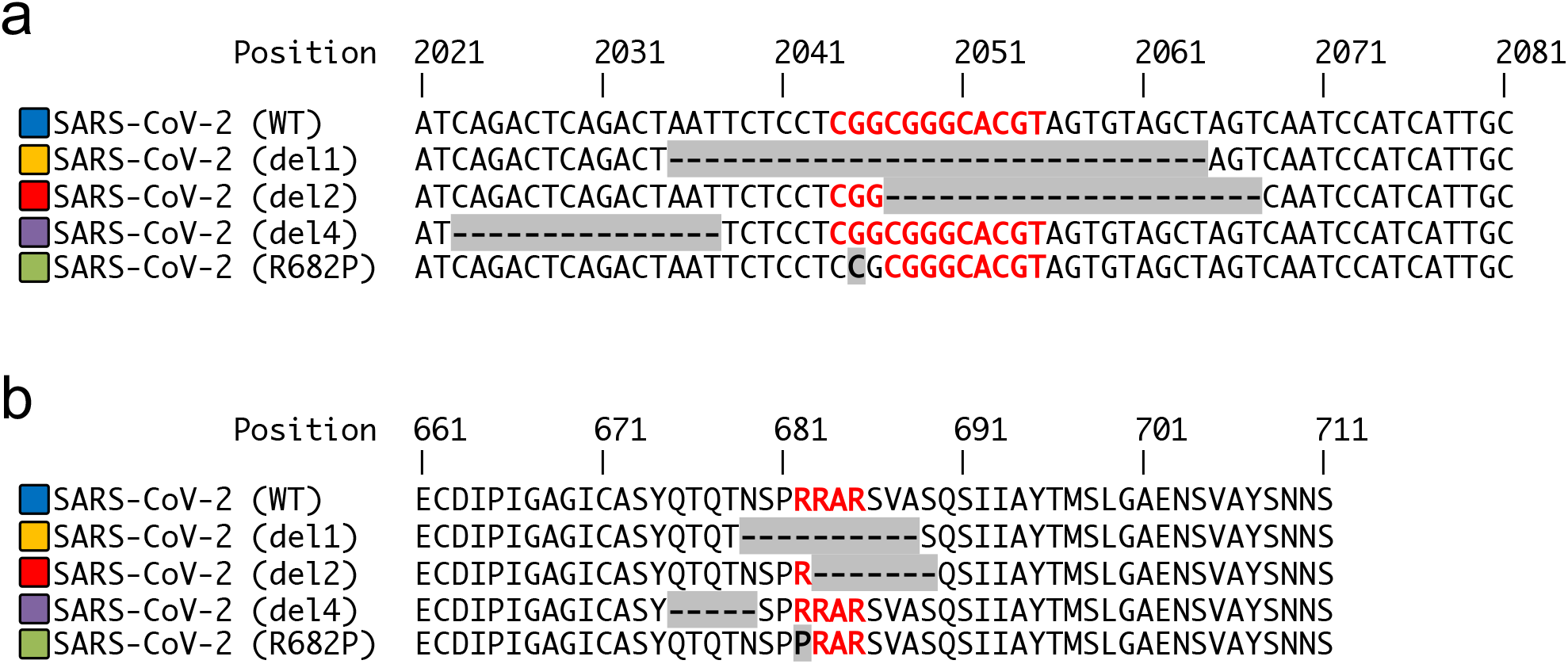
Multiple sequence alignment of S1/S2 cleavage site of wild type and SARS-CoV-2 variants. Multiple (a) nucleotide and (b) amino acid sequence alignments were constructed based on the sequence of WT and SARS-CoV-2 variants identified by deep-sequencing (related to Fig. 4). Infectious viruses of del4 and R682P were not isolated in this study. Nucleotide substitutions and deletions are shown as gray boxes. Sequence encoding the polybasic cleavage motif (RARR) at the S1/S2 cleavage site is highlighted in red.

**Fig. S3.**
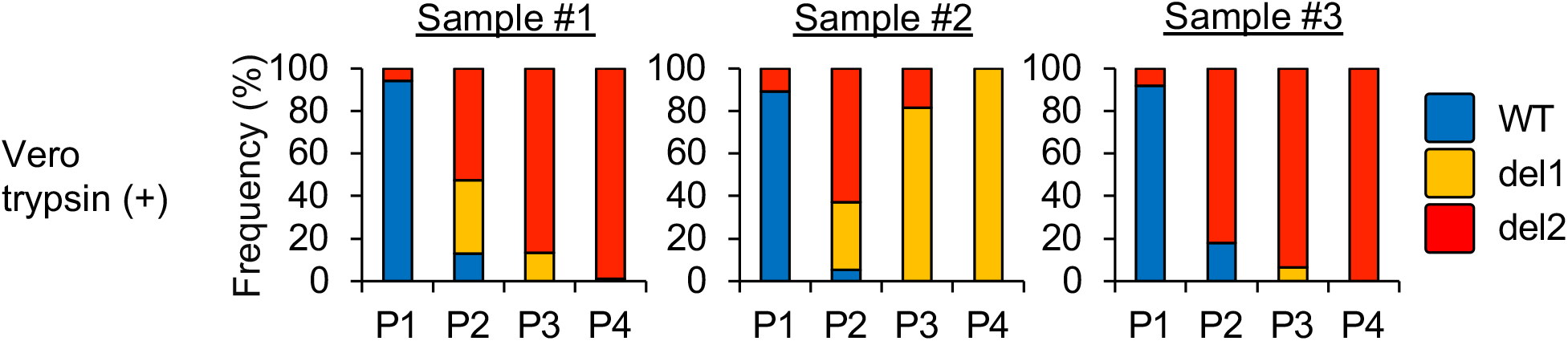
Frequencies of S gene mutants detected during SARS-CoV-2 propagation in Vero cells in the presence of trypsin. SARS-CoV-2 was serially passaged in Vero cells in serum free DMEM containing trypsin with three biological replicates. Nucleotide sequence diversity at viral S1/S2 cleavage site was determined by deep-sequencing.

## References

1. (WHO) WHO. Coronavirus disease (COVID-19) pandemic 2020 [cited 2020 6th July]. Available from: https://www.who.int/emergencies/diseases/novel-coronavirus-2019.

2. Naqvi AAT, Fatima K, Mohammad T, Fatima U, Singh IK, Singh A, et al. Insights into SARS-CoV-2 genome, structure, evolution, pathogenesis and therapies: Structural genomics approach. Biochim Biophys Acta Mol Basis Dis. 2020;1866(10):165878. Epub 2020/06/13. doi: 10.1016/j.bbadis.2020.165878. PubMed PMID: 32544429; PubMed Central PMCID: PMCPMC7293463.

3. Wang YT, Landeras-Bueno S, Hsieh LE, Terada Y, Kim K, Ley K, et al. Spiking Pandemic Potential: Structural and Immunological Aspects of SARS-CoV-2. Trends Microbiol. 2020. Epub 2020/05/20. doi: 10.1016/j.tim.2020.05.012. PubMed PMID: 32507543; PubMed Central PMCID: PMCPMC7237910.

4. Walls AC, Park YJ, Tortorici MA, Wall A, McGuire AT, Veesler D. Structure, Function, and Antigenicity of the SARS-CoV-2 Spike Glycoprotein. Cell. 2020;181(2):281–92.e6. Epub 2020/03/09. doi: 10.1016/j.cell.2020.02.058. PubMed PMID: 32155444; PubMed Central PMCID: PMCPMC7102599.

5. Wrapp D, Wang N, Corbett KS, Goldsmith JA, Hsieh CL, Abiona O, et al. Cryo-EM structure of the 2019-nCoV spike in the prefusion conformation. Science. 2020;367(6483):1260–3. Epub 2020/02/19. doi: 10.1126/science.abb2507. PubMed PMID: 32075877; PubMed Central PMCID: PMCPMC7164637.

6. Hoffmann M, Kleine-Weber H, Schroeder S, Krüger N, Herrler T, Erichsen S, et al. SARS-CoV-2 Cell Entry Depends on ACE2 and TMPRSS2 and Is Blocked by a Clinically Proven Protease Inhibitor. Cell. 2020;181(2):271–80.e8. Epub 2020/03/05. doi: 10.1016/j.cell.2020.02.052. PubMed PMID: 32142651; PubMed Central PMCID: PMCPMC7102627.

7. Hoffmann M, Kleine-Weber H, Pöhlmann S. A Multibasic Cleavage Site in the Spike Protein of SARS-CoV-2 Is Essential for Infection of Human Lung Cells. Mol Cell. 2020;78(4):779–84.e5. Epub 2020/05/01. doi: 10.1016/j.molcel.2020.04.022. PubMed PMID: 32362314; PubMed Central PMCID: PMCPMC7194065.

8. Andersen KG, Rambaut A, Lipkin WI, Holmes EC, Garry RF. The proximal origin of SARS-CoV-2. Nat Med. 2020;26(4):450–2. doi: 10.1038/s41591-020-0820-9. PubMed PMID: 32284615; PubMed Central PMCID: PMCPMC7095063.

9. Ren X, Glende J, Al-Falah M, de Vries V, Schwegmann-Wessels C, Qu X, et al. Analysis of ACE2 in polarized epithelial cells: surface expression and function as receptor for severe acute respiratory syndrome-associated coronavirus. J Gen Virol. 2006;87(Pt 6):1691–5. doi: 10.1099/vir.0.81749-0. PubMed PMID: 16690935.

10. Chu H, Chan JF-W, Yuen TT-T, Shuai H, Yuan S, Wang Y, et al. Comparative tropism, replication kinetics, and cell damage profiling of SARS-CoV-2 and SARS-CoV with implications for clinical manifestations, transmissibility, and laboratory studies of COVID-19: an observational study. The Lancet Microbe. 2020.

11. Matsuyama S, Nao N, Shirato K, Kawase M, Saito S, Takayama I, et al. Enhanced isolation of SARS-CoV-2 by TMPRSS2-expressing cells. Proc Natl Acad Sci U S A. 2020;117(13):7001–3. Epub 2020/03/12. doi: 10.1073/pnas.2002589117. PubMed PMID: 32165541; PubMed Central PMCID: PMCPMC7132130.

12. Liao K, Sikkema D, Wang C, Lee TN. Development of an enzymatic assay for the detection of neutralizing antibodies against therapeutic angiotensin-converting enzyme 2 (ACE2). J Immunol Methods. 2013;389(1-2):52–60. Epub 2013/01/05. doi: 10.1016/j.jim.2012.12.010. PubMed PMID: 23298658.

13. Böttcher-Friebertshäuser E, Stein DA, Klenk HD, Garten W. Inhibition of influenza virus infection in human airway cell cultures by an antisense peptide-conjugated morpholino oligomer targeting the hemagglutinin-activating protease TMPRSS2. J Virol. 2011;85(4):1554–62. Epub 2010/12/01. doi: 10.1128/JVI.01294-10. PubMed PMID: 21123387; PubMed Central PMCID: PMCPMC3028871.

14. Bertram S, Glowacka I, Blazejewska P, Soilleux E, Allen P, Danisch S, et al. TMPRSS2 and TMPRSS4 facilitate trypsin-independent spread of influenza virus in Caco-2 cells. J Virol. 2010;84(19):10016–25. Epub 2010/07/14. doi: 10.1128/JVI.00239-10. PubMed PMID: 20631123; PubMed Central PMCID: PMCPMC2937781.

15. Tai W, He L, Zhang X, Pu J, Voronin D, Jiang S, et al. Characterization of the receptor-binding domain (RBD) of 2019 novel coronavirus: implication for development of RBD protein as a viral attachment inhibitor and vaccine. Cell Mol Immunol. 2020;17(6):613–20. Epub 2020/03/19. doi: 10.1038/s41423-020-0400-4. PubMed PMID: 32203189; PubMed Central PMCID: PMCPMC7091888.

16. Kawase M, Shirato K, van der Hoek L, Taguchi F, Matsuyama S. Simultaneous treatment of human bronchial epithelial cells with serine and cysteine protease inhibitors prevents severe acute respiratory syndrome coronavirus entry. J Virol. 2012;86(12):6537–45. Epub 2012/04/11. doi: 10.1128/JVI.00094-12. PubMed PMID: 22496216; PubMed Central PMCID: PMCPMC3393535.

17. Scavone C, Brusco S, Bertini M, Sportiello L, Rafaniello C, Zoccoli A, et al. Current pharmacological treatments for COVID-19: What’s next? Br J Pharmacol. 2020. Epub 2020/04/24. doi: 10.1111/bph.15072. PubMed PMID: 32329520; PubMed Central PMCID: PMCPMC7264618.

18. Hoffmann M, Schroeder S, Kleine-Weber H, Müller MA, Drosten C, Pöhlmann S. Nafamostat Mesylate Blocks Activation of SARS-CoV-2: New Treatment Option for COVID-19. Antimicrob Agents Chemother. 2020;64(6). Epub 2020/05/21. doi: 10.1128/AAC.00754-20. PubMed PMID: 32312781; PubMed Central PMCID: PMCPMC7269515.

19. Yamamoto M, Kiso M, Sakai-Tagawa Y, Iwatsuki-Horimoto K, Imai M, Takeda M, et al. The Anticoagulant Nafamostat Potently Inhibits SARS-CoV-2 S Protein-Mediated Fusion in a Cell Fusion Assay System and Viral Infection In Vitro in a Cell-Type-Dependent Manner. Viruses. 2020;12(6). Epub 2020/06/10. doi: 10.3390/v12060629. PubMed PMID: 32532094.

20. Ogando NS, Dalebout TJ, Zevenhoven-Dobbe JC, Limpens RWAL, van der Meer Y, Caly L, et al. SARS-coronavirus-2 replication in Vero E6 cells: replication kinetics, rapid adaptation and cytopathology. J Gen Virol. 2020. Epub 2020/06/22. doi: 10.1099/jgv.0.001453. PubMed PMID: 32568027.

21. Wang M, Cao R, Zhang L, Yang X, Liu J, Xu M, et al. Remdesivir and chloroquine effectively inhibit the recently emerged novel coronavirus (2019-nCoV) in vitro. Cell Res. 2020;30(3):269–71. Epub 2020/02/04. doi: 10.1038/s41422-020-0282-0. PubMed PMID: 32020029; PubMed Central PMCID: PMCPMC7054408.

22. Wada Y, Orba Y, Sasaki M, Kobayashi S, Carr MJ, Nobori H, et al. Discovery of a novel antiviral agent targeting the nonstructural protein 4 (nsP4) of chikungunya virus. Virology. 2017;505:102–12. Epub 2017/02/23. doi: 10.1016/j.virol.2017.02.014. PubMed PMID: 28236746.

23. Shirogane Y, Takeda M, Iwasaki M, Ishiguro N, Takeuchi H, Nakatsu Y, et al. Efficient multiplication of human metapneumovirus in Vero cells expressing the transmembrane serine protease TMPRSS2. J Virol. 2008;82(17):8942–6. Epub 2008/06/18. doi: 10.1128/JVI.00676-08. PubMed PMID: 18562527; PubMed Central PMCID: PMCPMC2519639.

24. Ou X, Liu Y, Lei X, Li P, Mi D, Ren L, et al. Characterization of spike glycoprotein of SARS-CoV-2 on virus entry and its immune cross-reactivity with SARS-CoV. Nat Commun. 2020;11(1):1620. Epub 2020/03/27. doi: 10.1038/s41467-020-15562-9. PubMed PMID: 32221306; PubMed Central PMCID: PMCPMC7100515.

25. Lau SY, Wang P, Mok BW, Zhang AJ, Chu H, Lee AC, et al. Attenuated SARS-CoV-2 variants with deletions at the S1/S2 junction. Emerg Microbes Infect. 2020;9(1):837–42. doi: 10.1080/22221751.2020.1756700. PubMed PMID: 32301390; PubMed Central PMCID: PMCPMC7241555.

26. Davidson AD, Williamson MK, Lewis S, Shoemark D, Carroll MW, Heesom KJ, et al. Characterisation of the transcriptome and proteome of SARS-CoV-2 reveals a cell passage induced in-frame deletion of the furin-like cleavage site from the spike glycoprotein. Genome Med. 2020;12(1):68. Epub 2020/07/28. doi: 10.1186/s13073-020-00763-0. PubMed PMID: 32723359; PubMed Central PMCID: PMCPMC7386171.

27. Corman VM, Landt O, Kaiser M, Molenkamp R, Meijer A, Chu DK, et al. Detection of 2019 novel coronavirus (2019-nCoV) by real-time RT-PCR. Euro Surveill. 2020;25(3). doi: 10.2807/1560-7917.ES.2020.25.3.2000045. PubMed PMID: 31992387; PubMed Central PMCID: PMCPMC6988269.

28. Overbergh L, Kyama CM, Valckx D, Debrock S, Mwenda JM, Mathieu C, et al. Validation of real-time RT-PCR assays for mRNA quantification in baboons. Cytokine. 2005;31(6):454–8. doi: 10.1016/j.cyto.2005.07.002. PubMed PMID: 16129617.

29. Pauwels R, Balzarini J, Baba M, Snoeck R, Schols D, Herdewijn P, et al. Rapid and automated tetrazolium-based colorimetric assay for the detection of anti-HIV compounds. J Virol Methods. 1988;20(4):309–21. doi: 10.1016/0166-0934(88)90134-6. PubMed PMID: 2460479.

